# Ethylene Insensitive 3-Like 2 (EIL2) is a *Brassicaceae*-specific transcriptional regulator involved in fine-tuning ethylene-dependent hypocotyl elongation, lateral root formation and flowering time in *Arabidopsis thaliana*

**DOI:** 10.1101/2022.02.09.479754

**Authors:** Maarten Houben, John Vaughan-Hirsch, Wangshu Mou, Bram Van de Poel

**Affiliations:** Division of Crop Biotechnics, Department of Biosystems, University of Leuven, Willem de Croylaan 42, 3001 Leuven, Belgium

**Keywords:** Transcription factor, Ethylene, *Brassicaceae*, lateral roots, hypocotyl, flowering

## Abstract

Ethylene signaling directs a pleiotropy of developmental processes in plants. In Arabidopsis, ethylene signaling converges at the master transcription factor Ethylene Insensitive 3 (EIN3), which has five homologs, EIN3-like 1-5 (EIL1-5). EIL1 is most characterized and operates similar as EIN3, while EIL3-5 are not involved in ethylene signaling. EIL2 remains less investigated. Our phylogenetic analysis revealed that EIL2 homologs are only retrieved in the *Brassicaceae* family, suggesting EIL2 diverged to have specific functions in the mustard family. By characterizing *eil2* mutants, we found that EIL2 is involved in regulating ethylene-specific developmental processes in *Arabidopsis thaliana*, albeit in a more subtle way compared to EIN3/EIL1. EIL2 steers ethylene-triggered hypocotyl elongation in light-grown seedlings and it is involved in lateral root formation. Furthermore, EIL2 takes part in regulating flowering time as *eil2* mutants flower on average one day earlier and have fewer leaves. A pEIL2:EIL2:GFP translational reporter line revealed that EIL2 protein abundance is restricted to the stele of young developing roots. *EIL2* expression, and not EIL2 protein stability, is regulated by ethylene in an EIN3/EIL1-dependent way. Despite EIL2 taking part in several developmental processes, the precise upstream and downstream regulation of this ethylene- and *Brassicaceae*-specific transcription factor remains to be elucidated.

**Highlight:** EIL2 is an ethylene-dependent transcription factor that evolved exclusively in the mustard family and operates downstream from EIN3/EIL1. *EIL2* is involved in fine-tuning hypocotyl elongation, lateral root formation and flowering time in *Arabidopsis thaliana*.

## Introduction

The volatile plant hormone ethylene regulates many developmental processes and is involved in several biotic and abiotic stress responses in plants^1,2^. During the last decades, the ethylene signal transduction pathway has been elucidated, mainly using *Arabidopsis thaliana* as a model species. However, the ethylene signaling pathway is much more ancient, and has likely been established in ancestral charophyte green algae^3^, and seems highly conserved among embryophytes^4,5^. Ethylene signaling starts with the perception of ethylene gas at the endoplasmic reticulum membrane^6,7^. Ethylene sensing is mediated by a group of five ethylene receptors (ETR1, ETR2, ERS1, ERS2 & EIN4) in Arabidopsis, which are negative regulators of ethylene responses^8^. The receptors belong to the family of bacterial derived two-component signal transduction system^9^, and have an ER localized N-terminal transmembrane ethylene binding domain that senses ethylene gas, a cytosolic GAF domain and a C-terminal histidine-kinase domain^10^. The C-terminal part of the ethylene receptors interact with a downstream cytosolic kinase Constitutive Triple Response 1 (CTR1)^11^, which is also a negative regulator of ethylene signaling^12^. In the absence of ethylene, CTR1 phosphorylates Ethylene Insensitive 2 (EIN2)^13^, which is a positive regulator of ethylene signaling and has an ER bound N-terminal transmembrane domain (with protein similarity to Nramp metal transporters) and a cytosolic C-terminal domain^14,15^. When EIN2 is phosphorylated by CTR1, EIN2 is degraded via two F-box EIN2 Targeting Proteins (ETP1/2)^16^ and fails to induce downstream ethylene signaling. When ethylene binds to the receptor, CTR1 is inactivated and consequently EIN2 phosphorylation is prevented, leading to EIN2 cleavage and the migration of its C-terminal end (CEND) to the nucleus^13,17,18^. Subsequently, the EIN2 CEND will activate Ethylene Insensitive 3 (EIN3)^15^ via an unknown mechanism. In the absence of ethylene, EIN3 is turned over by the 26S proteasome after ubiquitination by the F-box protein EIN3 Binding Factors 1 and 2 (EBF1/2)^19–21^. The translation efficiency of these EBFs is also directly inhibited by the EIN2 CEND in P-bodies^22,23^. Upon ethylene signaling, EIN3 is stabilized and activates Ethylene Responsive transcription Factors (ERFs) which subsequently activate secondary target genes leading to ethylene responses^24^. EIN3 binds to Ethylene Binding Sites (EBS), in the promotor sequences of downstream ERFs^24,25^. In addition to this canonical signaling, the EIN2 CEND also stimulates acetylation of histones H3K14 and H3K23 via the interaction with EIN2 Nuclear Associated Protein1 (ENAP1) to facilitate ethylene-depended gene expression^26,27^. ENAP1 also interacts with EIN3, the master transcription factor that has five homologs EIN3-like1 – 5 (EIL1-5)^28^.

Alongside EIN3, EIL1 and EIL2 were shown to take part in ethylene signaling. This was demonstrated by constitutive expression (*35S*) of both *EIL1* and *EIL2* in the *ein3-1* mutant, which was sufficient to partially rescue its ethylene insensitive phenotype^28^. However, the very strong ethylene insensitivity of the *ein3eil1* double and *ein3eil1ebf1ebf2* quadruple mutants suggested that EIN3 and EIL1 are able to process the majority of the ethylene signal, independent of the other EIL’s^29^. Consequently, little is known about the exact role of the EIL2-5 homologs. *EIL2* has a much more restricted gene expression pattern compared to *EIL1*, with only a weak expression in the root tip and pollen^30^. While EIL2 is still involved in ethylene signaling^28^, EIL3-5 have probably other functions independent of ethylene. EIL3, also called SLIM1 (Sulfur Limitation 1), has not been linked to ethylene signaling, and is the only EIL known to be involved in regulating sulfur deficiency^31,32^. The roles of EIL4 and EIL5 remain unknown, and both genes have a very low expression level, and are mainly expressed in developing dark-grown seedlings^30^.

In this study, we zoom in on the function of EIL2 and its role in ethylene signaling to regulate plant development. We have found that EIL2 evolved as a *Brassicaceae*-specific transcription factor, and that *eil2* mutants shows subtle vegetative and generative phenotypes, yet in an ethylene-dependent way and regulated by EIN3/EIL1.

## Materials and methods

### Plant material, growth conditions and phenotyping

*Arabidopsis thaliana* ecotype Col-0 was used as wild-type control plants. The *eil2* (SALK_202725C) and *ein3-1* (N8052) T-DNA insertion mutants were obtained from the Nottingham Arabidopsis Stock Centre (NASC) and *ein3-1eil1-1* was a gift from Dr. Caren Chang (University of Maryland, USA). Seeds were surface sterilized by incubation in a 90 % ethanol solution (v/v) for one minute, followed by a one minute incubation in a 70 % ethanol solution (v/v) and a one minute incubation in a 2 % bleach solution (NaOCl). The seeds were then washed in sterile H_2_O and imbibed at 4 °C for three days. Next, seeds were sown on Petri dishes containing 0.5 x Murashige and Skoog (MS) basal salt medium with 0.8 % agar (pH 5.7), supplemented with 1-aminocyclopropane-1-carboxylic acid (ACC) concentrations (final ACC concentrations 0 μM, 0.1 μM, 0.2 μM, 0.5 μM, 1 μM, 2 μM and 5 μM).

For triple response phenotyping, petri dishes containing seeds were illuminated for approximately three hours at room temperature with red light to promote germination and subsequently incubated in the dark at 21 °C for 4 days. Afterwards, seedlings were photographed to measure hypocotyl and root length. For light phenotyping, petri dishes containing seeds were grown vertically in an *in vitro* growth room (16 h light – 8 h dark; 100 μmol.m^−2^.s^−1^; 21 °C). Ten days after sowing, pictures were taken to analyze hypocotyl and root length. Image analysis was done with Imagej.

For phenotyping mature plants, WT and e/72 plants were grown in the same conditions in a controlled growth chamber (16 h light - 8 h dark; 110 μmol.m^−2^.s^−1^; 21 °C). Flowering time was assessed by counting the number of days until bolting. The leaf number and area at bolting was recorded by cutting all rosette leaves and arranging them in order of age before imaging with a Nikon D3300 camera. Total leaf area was calculated using ImageJ. Flower and silique morphology analysis was done by imaging flowers at stage 14^33^ or mature green siliques (7 to 10 days post pollination) from the middle of the primary inflorescence with a stereomicroscope (Olympus SZX). For siliques, the carpel was removed using a 27.5-gauge needle prior to imaging. Healthy fertilized embryos were counted as a seed for each silique, and silique length was measured using ImageJ.

### Ethylene measurements

Ethylene production rates of 4-day old dark-grown seedlings (21 °C) were quantified in airtight GC-vials on MS medium supplemented with 2 μM ACC. One mL of headspace was sampled with a 1 mL syringe and injected into a Shimadzu gas chromatograph (GC-2014). The GC was equipped with a Porapak R50/80 stainless steel column (3 m x 3 mm) with N_2_ as a carrier gas (35 mL/min). Both the injector and column temperature was set at 150 °C. Ethylene was detected using a flame ionization detector at a temperature of 250 °C.

### Genome wide association mapping (GWAS)

GWAS was conducted using ethylene production data originating from 4-day old dark-grown seedlings grown on 2 μM ACC, for 155 different Arabidopsis ecotypes (5 biological repeats per ecotype). The ethylene production was normalized per germinated seedling, and the data was analyzed using the online GWAPP portal^34^. All phenotypic data were analyzed using both a linear regression model and the accelerated linear mixed model in combination with or without a Box-Cox transformation. These four analysis methods were compared and the minor allele frequency (MAF) threshold was set at 10 % and the −log(P) value was set to be minimum 5. A list of possible associated SNPs was manually verified and curated based on SNP identity and loci. Manhattan plots were generated using the GWAPP portal.

### Phylogenetic analysis of the *EIN3/EIL* gene family

BLASTp jobs were done for 20 plant species using the Phytozome (v12.1.) database and BLASTp on the NCBI site (https://blast.ncbi.nlm.nih.gov) using the *Arabidopsis thaliana* EIN3 (AT3G20770) protein sequence as query. Top BLASTp hits were only retained after a positive reciprocal BLAST search. Protein sequences were aligned in Geneious Prime^®^ (v2020.2.3) using the MUSCLE alignment plugin. The phylogenetic tree was constructed using RAxML (v8.2.11) for best-scoring maximum likelihood tree with rapid bootstrapping (1000 bootstrap replicates). iTOL was used for the display and annotation of the tree^35^.

### *In vitro* and *in vivo* pollen tube germination and growth assays

The *in vitro* pollen tube growth assay was performed as described by Boavida et al. (2007)^36^. Dehiscent anthers were tapped to spread pollen grains on a thin layer of agar pollen germination medium (8 % sucrose; 0.01 % Boric acid; 1 mM CaCl_2_; 1 mM Ca(NO_3_)_2_; 1 mM MgSO_4_; 0.5 % Noble agar (Difco)) prepared in a glass bottom petri dish (35 mm diameter). After incubation in a humidity chamber for 3.5 h, pollen tubes were imaged using the Leica Thunder fluorescence microscope and analyzed with ImageJ.

For *in vivo* pollen tube growth, aniline blue staining of pollen tubes in pistils was performed according to Mori et al. (2006)^37^. Flowers were pre-emasculated at stage 12b and pollinated 24 h later. Subsequently, 6.5 h after pollination, the pistils were incubated in fixation solution containing ethanol:acetic acid (3:1) for 2 h at room temperature and rehydrated in ethanol (steps of 20 % each 10 min). Next, the pistils were softened in 8 M NaOH overnight and washed with distilled water for 10 min. The pistils were then stained overnight with decolorized aniline blue 0.1 % (w/v) in the dark, and pollen tubes were observed using a Leica Thunder fluorescence microscope with a 20 × objective.

### Plasmid construction and Arabidopsis transformation

Genomic DNA extracted from Col-0 wild-type seedlings was used as a template to amply the *EIL2* coding sequence and the *EIL2* promotor. All primers used in this study are listed in Supplementary Table S1. For the creation of the native translational reporter line *pEIL2:EIL2:GFP*, the promoter region 2632 bases upstream from the ATG start codon of *EIL2* was amplified together with the *EIL2* CDS. The amplified fragment was cloned into Ncol/Kpnl-digested pCambia1302 carrying a GFP, using Gibson assembly. The three resulting constructs were transformed into *Arabidopsis thaliana* Col-0 wild-type using the floral dip method, and homozygous T3 lines selected for use in experiments.

### *pEIL2:EIL2:GFP* imaging

Seeds of the *pEIL2:EIL2:GFP* reporter line were sterilized as described above and grown for 7 days in the same conditions as described above. Seedlings were then incubated for 4 h in ½ strength MS liquid media supplemented with 10 μM ACC, 50 μM MG132, or both, alongside a control with an equal volume of DMSO added. Col-0 was also incubated with DMSO as a control for imaging. After incubation, seedling root tips were immediately imaged using a Leica Thunder fluorescence microscope with a GFP filter cube (excitation 450-490 nm; emission 500-550 nm).

### RT-qPCR analysis

For RNA extractions, seedlings or flowers were harvested, crushed in liquid nitrogen, and RNA extracted using the GeneJET Plant RNA extraction kit (Thermo). Following RNA extraction, DNA was removed using the RapidOut DNA removal kit (Thermo) and RNA quantified using a Nanodrop 2000 (Nanodrop Technology). cDNA was then prepared from 100 ng RNA using the iScript cDNA synthesis kit (Bio-Rad). qRT-PCR reactions were set up with 4 biological replicates, using Sso Advanced Universal SYBR Green Supermix (Bio-Rad) and 1 ng cDNA, and run in a Bio-Rad CFX96 qPCR machine with the following cycling conditions: 98 °C for 30 sec (initial denaturation) followed by 40 cycles of 98 °C for 15 sec (denaturation), and 60 °C for 30 sec (annealing and extension), with a final melt curve analysis performed at the end of the run to check for single amplicons. Two housekeeping genes were analyzed (*UBC21* and *UBQ10*) for normalization and relative expression calculated using the delta-delta Ct method.

### Statistical analysis

The results were processed using ‘Rstudio’ (version 3.2.5, Boston, USA). The Shapiro-Wilk test (‘shapiro.test’) was used to test the normal distribution of the data. The function (‘boxplot’) was used to identify possible outliers, which were defined as data points outside 1.5 times the interquartile range above the upper quartile or below the lower quartile. The “Agricolae” package was used for the multiple comparisons of treatments by means of Tukey (‘HSD.test’) when the assumption of normality was met. When the assumption of normality was not met, the non-parametric Kruskal–Wallis one-way analysis of variances test was used (‘kruskal’). All test were done with a 95 % confidence interval (α = 0.05).

## Results

We performed a quantitative analysis of ethylene production on 4-day old dark-grown Arabidopsis seedlings of 155 different ecotypes, and used this dataset to perform a Genome Wide Association (GWA) analysis. Through this GWAS experiment, we were able to identify a highly associated SNP on chromosome 5, linked to the *EIL2* locus (AT5G21120), more specifically to the *EIL2* promotor region 1223 base pairs upstream of the start-codon (Figure 1A). This urged us to study the involvement of EIL2 in regulating ethylene biosynthesis and signaling.

**Figure 1:**
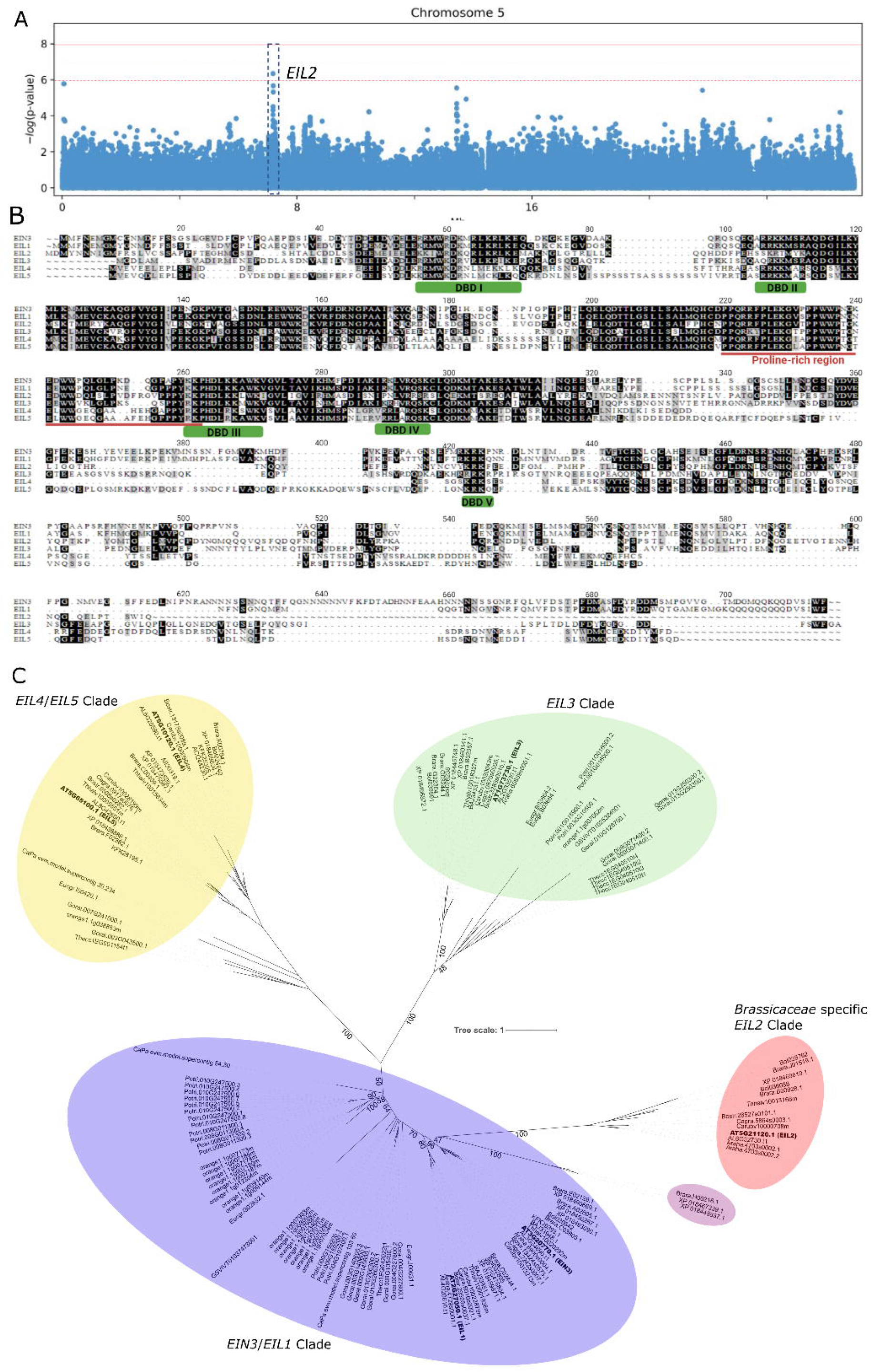
EIL2 identification and phylogenetic analysis. (A) Manhattan plot of chromosome 5 of *Arabidopsis thaliana* showing the *EIL2* locus significantly associated with ethylene production as the result of a GWAS analysis of 4-day old dark-grown seedlings fed with 2 μM ACC. (B) EIL2 protein alignment with its closest homologs EIN3, EIL1 and EIL3-5. Green bars indicate conserved DNA binding domains (DBD1-5) and the red line indicates a conserved proline rich region. (C) Maximal likelihood phylogenetic tree of EIL2 and its homologs (EIN3, EIL1, EIL3-5) of angiosperms including *Arabidopsis thaliana, Arabidopsis halleri, Arabidopsis lyrata, Arabis alpine, Boechera stricta, Brassica oleracea capitate, Brassica rapa, Capsella grandiflora, Capsella rubella, Cardamine hirsute, Carica papaya, Citrus sinensis, Eucalyptus grandis, Eutrema halophilum, Eutrema salsugineum, Gossypium raimondii, Populus trichocarpa, Raphanus sativus, Theobroma cacao* and *Vitis vinifera*. The different EIL clades are colored based on phylogenetic grouping: EIN3/EIL1 homologs (purple), EIL2 homologs (red; only *Brassicaceae*), EIL3 homologs (green) and EIL4-5 homologs (yellow). Bootstrap values (1000 replicates) for the main branches are depicted on the tree.

### *EIL2* codes for a specialized *Brassicaceae*-specific homolog of EIN3

First, we performed a phylogenetic analysis to study EIL2’s relation with the master transcription factor EIN3. Therefore we aligned the EIN3 protein sequence with its closest homologs EIL1 – EIL5 (Figure 1B), which showed that both EIL1-5 are missing a C-terminal asparagine rich region, and that EIL2 completely lacks a large C-terminal part. EIL2 showed only 44.7 % similarity with EIN3 and 46.1 % similarity with EIL1 (see Supplementary Table S2). Interestingly, EIL2 also showed sequence similarity with non-ethylene specific EIL3 (36.7 %), EIL4 (36.4 %) and EIL5 (36.3 %) homologs (Supplementary Table S2), which also possess the conserved DNA-binding motif present in EIN3 (Figure 1B). The major differences between EIN3/EIL1-2 and EIL3-5 is that the latter miss about 20 amino acids in a non-conserved region (Figure 1B) and a 10 amino acid long N-terminal tail that contains a putative N-myristoylation signal peptide.

A broader phylogenetic analysis of all EIN3 and EIL homologs from 24 different plant species ranging from angiosperms to green algae (Supplementary Figure S1) revealed that EIL2 branches out as a small sub-clade from the larger EIN3/EIL1 clade. This EIN3/EIL1 clade is also phylogenetically very distant from the other Arabidopsis EIL3 and EIL4-5 groups, as well as a clade comprising ancestral EIN3-homologs from more-distant charophyte algae (*Spirogyra pratensis*) and early diverging land plants (*Marchantia polymorpha*, *Selaginella moellendorffii* and *Physcomitrella patens)*. The EIN3 and EIL1/EIL2 sub-clade clusters together only with EIN3 homologs from angiosperms, and not with EIN3 homologs from other species. Therefore, we believe that EIN3 and EIL1/EIL2 proteins evolved to become highly-specialized transcription factors, tailored to angiosperm-specific transcriptional control of ethylene responses. The divergence of this EIN3 sub-clade and the outbranching of EIL2 in this group, made us wonder about the specialization of EIL2. Therefore, we constructed a new phylogenetic tree, containing only angiosperms and additional *Brassicaceae* species (Figure 1C). Remarkably, the EIL2 homologs were only retrieved in *Brassicaceae* species, suggesting EIL2 diverged from an ancestral EIN3 protein to obtain a specialized function in the *Brassicaceae* family.

### EIL2 modulates subtle vegetative ethylene-dependent phenotypes

Based on Gene-investigator and eFP-browser^38^ expression datasets, *EIL2* seems to be predominantly expressed in the mature embryo, a young developing root and pollen grains. More specifically, *EIL2* shows the strongest expression in the root phloem and the two pollen sperm cells (Supplementary Figure S2 and S3). This expression profile urged us to examine the role of EIL2 in seedling ethylene production and in dark- and light-grown seedling development. We deployed an *eil2* T-DNA transgenic line (SALK_202725C; Figure 2A), which did not show any detectable *EIL2* expression in flowers (Figure 2B). Next, we analyzed ethylene production levels of dark-grown seedlings, but to our surprise, we did not observe any significant difference between Col-0 and *eil2* knockout mutants (Figure 2C), while *ein3* and *ein3eil1* seedlings produced significantly more ethylene. These finding are peculiar, and do not corroborate the GWAS results for which we measured ethylene production levels of dark-grown seedlings. Perhaps this indicates that a reduced *EIL2* gene expression does not associate with an altered ethylene production in the Col-0 ecotype. It is possible that EIL2 still influences ethylene production levels in other Arabidopsis ecotypes and that therefore we identified this gene in our GWA study.

**Figure 2:**
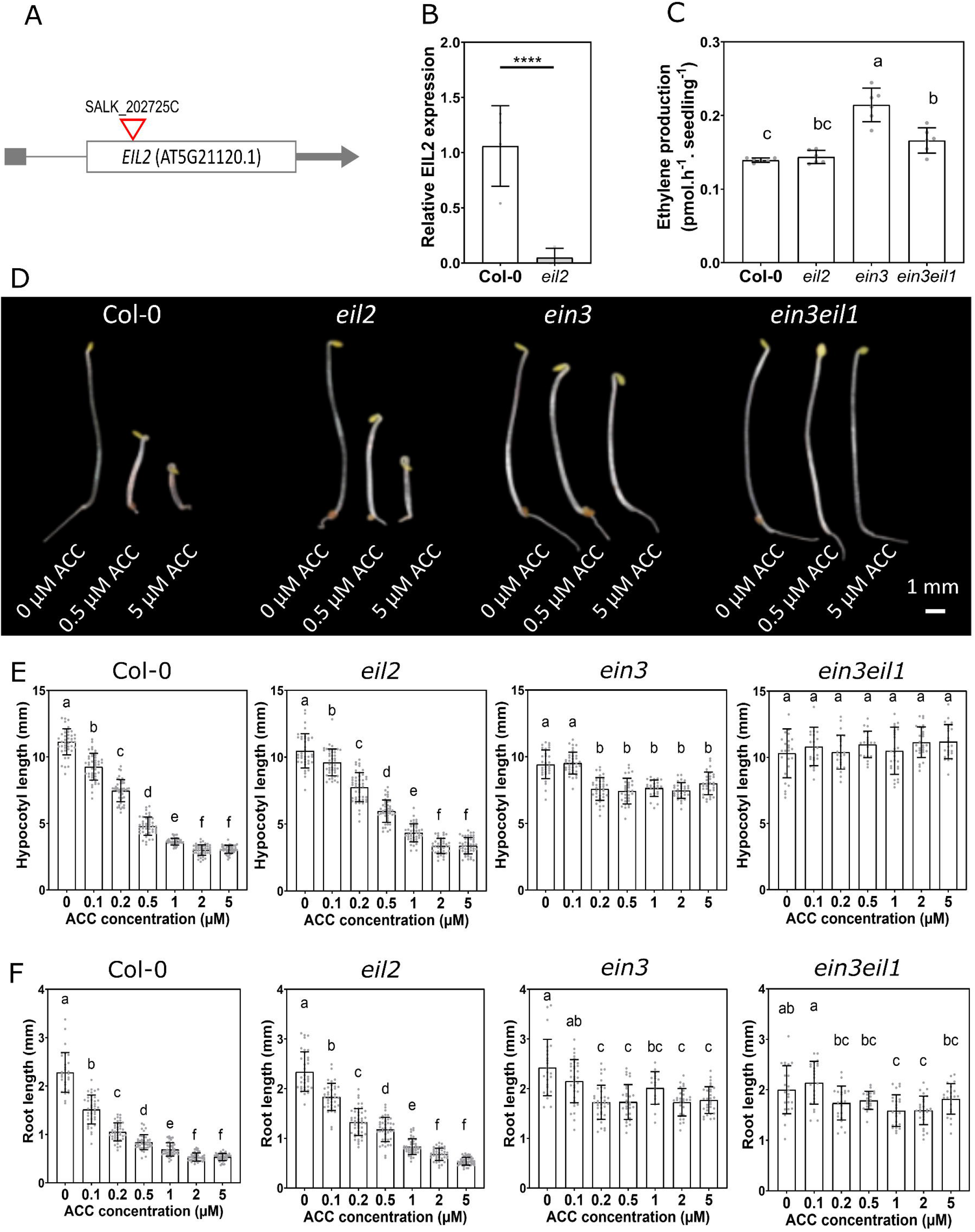
Subtle vegetative phenotypes of dark-grown *eil2* seedlings. (A) *EIL2* gene model with the T-DNA insert location of SALK line 202725C presented in red. (B) *EIL2* gene expression in wild type and the *eil2* flowers. (C) Ethylene production level of 4-day old dark-grown seedlings of Col-0, *eil2, ein3* and *ein3eil1* mutants supplemented with 2 μM ACC. (D) Representative seedlings, (E) hypocotyl length and (F) root length of 4-day old dark-grown seedlings of Col-0, *eil2*, *ein3*, *ein3eil1* treated with different doses of ACC (0 - 5 μM).

Next, we assessed the triple response phenotype for *eil2*, *ein3* and *ein3eil1* mutants (Figure 2D) using an ACC dose response curve as a substitute for ethylene gas. *Eil2* seedlings showed a mild ethylene-insensitivity in hypocotyl elongation towards intermediate ACC levels (0.5 −1 μM; Figure 2E), while *ein3* and *ein3eil1* hypocotyls showed a much stronger insensitivity towards ACC. The roots of dark-grown *eil2* mutants were slightly more insensitive towards ACC compared to Col-0 (Figure 2F). Again, the ethylene insensitive root phenotypes of *ein3* and *ein3eil1* seedlings were more pronounced.

For 10-day old light-grown seedlings, we observed that *eil2* plants showed a complete insensitivity towards the ACC-induced hypocotyl elongation at 5 μM ACC, in a similar way as *ein3* and *ein3eil1* mutants (Figure 3A & 3C). The growth of the main root of light-grown *eil2* plants did not show any insensitivity towards different concentrations of ACC, while *ein3* and *ein3eil1* mutants showed a partial and strong root insensitivity towards low and medium high ACC levels respectively (Figure 3B). The *eil2*mutant also showed an increased root hair growth upon ACC treatment, similar as for the wild type plants, but absent in the *ein3* and *sein3eil1* mutants, as was previously described^39^. Despite the normal sensitivity of the primary root of *eil2* plants towards ACC, *eil2* plants did show a drastic reduction in the number of lateral roots (Figure 3D), which was even more pronounced in the *ein3* and *ein3eil1* mutants.

**Figure 3.**
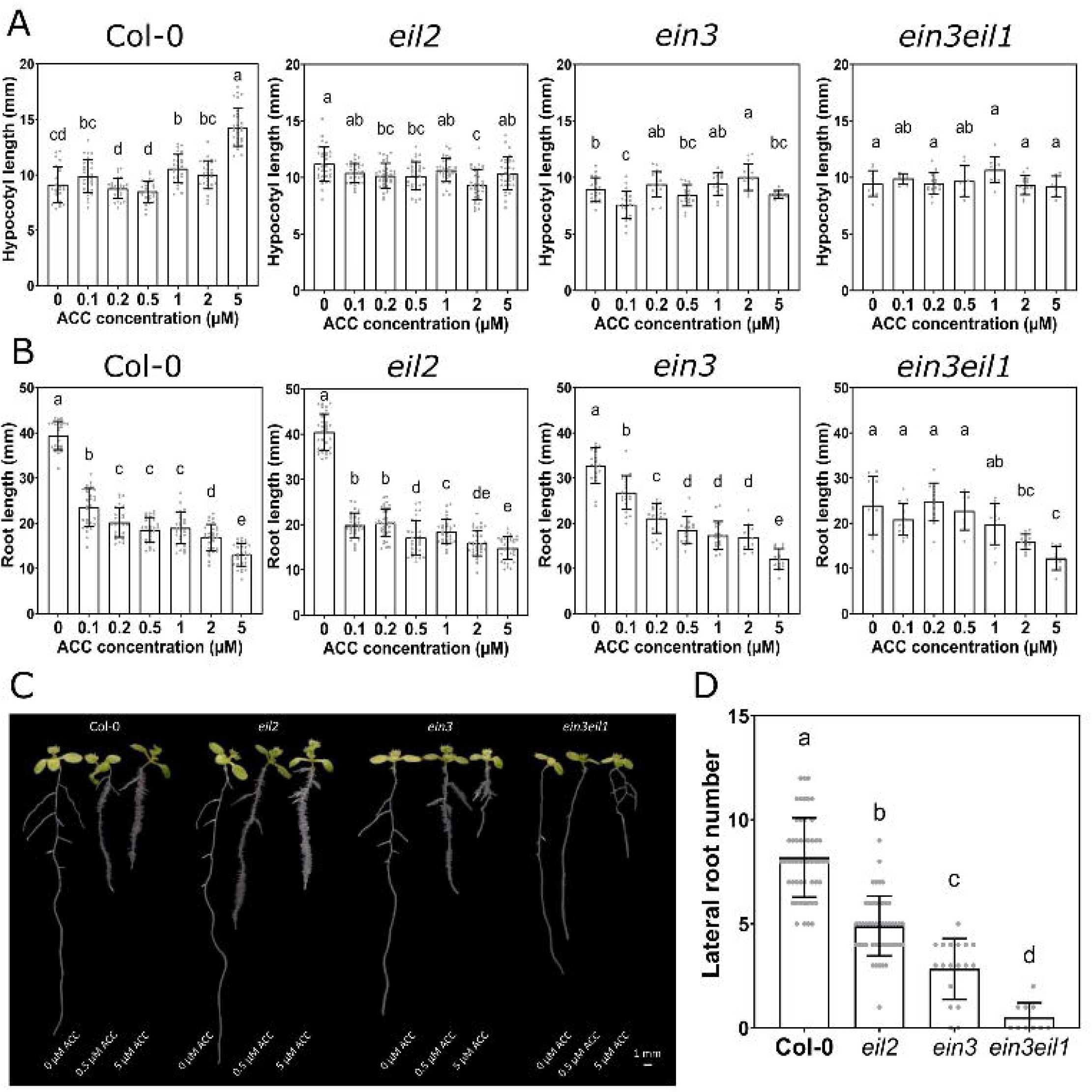
Subtle vegetative phenotypes of light-grown *eil2* seedlings. (A) Hypocotyl length and (B) root length of 10-day old light-grown seedlings of Col-0, *eil2*, *ein3*, *ein3eil1* treated with different doses of ACC (0 - 5 μM). (C) Representative seedlings of 10-day old light-grown seedlings of Col-0, *eil2, ein3* and *ein3eil1* treated with 0, 0.5 and 5 μM ACC. (D) Number of lateral roots of 10-day old light-grown seedlings of Col-0, *eil2 ein3* and *ein3eil1* in the absence of ACC.

### EIL2 mildly modulates flowering time and silique growth

Besides vegetative phenotypes, we also investigated if *eil2* plants show any generative phenotypes. We noticed that *eil2* plants consistently flowered about 1 day earlier compared to Col-0 (Figure 4A). This subtle early flowering phenotype of *eil2* was small but significant, and was corroborated by the observation that *eil2* plants had a significant lower total leaf area at bolting (Figure 4B & D). The reduced total leaf area was probably caused by a lower number of leaves developed until flowering time (Figure 4C-D), indicating that EIL2 is involved in leaf development.

**Figure 4.**
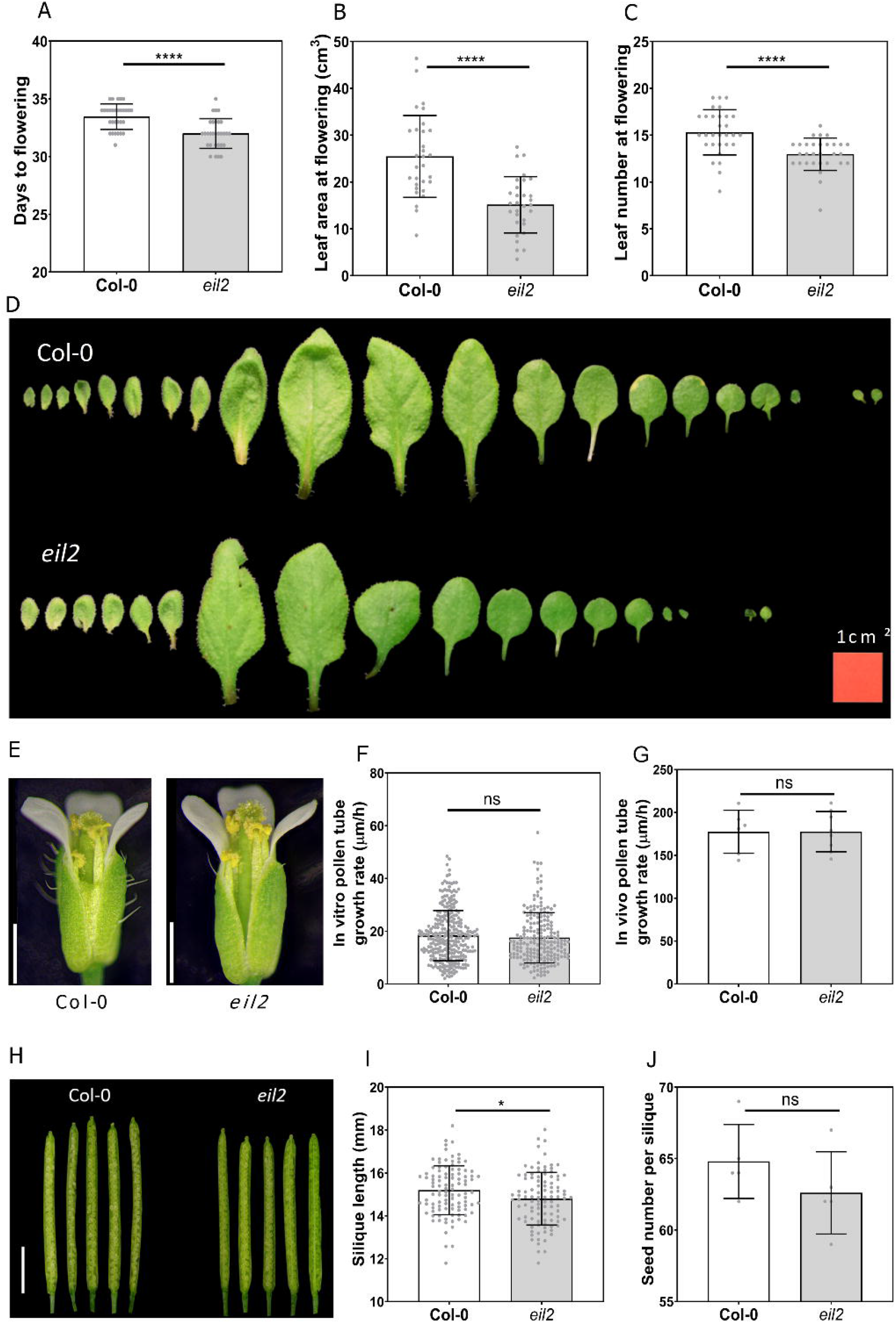
Generative and reproductive phenotypes of *eil2* plants. (A) *Eil2* plants flower on average 1 day earlier than Col-0 wild type plants. (B) Leaf area at flowering of Col-0 and *eil2*. (C) Leaf number at flowering of Col-0 and *eil2*. (D) Representative dissected rosettes of Col-0 and *eil2* plants at flowering. (E) Flower morphology of Col-0 and *eil2*. (F) *In vitro* and (G) *in vivo* pollen tube development of Col-0 and *eil2* pollen. (H) Representative siliques, (I) silique length and (J) seed number per silique of Col-0 and *eil2* plants.

Because microarray data indicates that *EIL2* has the highest expression in pollen sperm cells (Supplementary Figure S2), we examined if any reproductive or fertility defects were present in *eil2* plants. At first sight, flower morphology looks very similar to wild-type plants (Figure 4E), with no abnormal development of anthers, stigma, pistil, petals nor sepals. We also did not observe any defect in pollen tube development, nor under *in vitro* conditions (Figure 4F), nor under *in vivo* conditions (Figure 4G, Supplementary Figure S4 and S5). However, we did observe a small but significant reduction in the silique length of *eil2* plants, which was about 1 mm shorter (Figure 4H-I), indicating that EIL2 is mildly involved in silique growth. This did not manifest in a change in fertility as assessed by the total seed number per silique (Figure 4J).

### *EIL2* expression is weak but induced by ethylene and EIN3/EIL1 dependent

In order to examine the localization and stability of EIL2 *in planta*, we made a translational reporter line (C-terminal GFP) driven by the native *EIL2* promoter. In this line, an EIL2-GFP signal was only observed in the stele of the root when seedlings were treated with ACC (Figure 5A) or ethylene (Supplementary Figure S6). Since both ACC and ethylene treatment show a similar response, this rules out the possibility of ethylene-independent induction of EIL2 by ACC itself. To determine whether EIL2 is regulated at the level of protein stability, we imaged the EIL2-GFP signal after treatment with the proteasome inhibitor MG132 (Figure 5A). This gave no observable EIL2-GFP signal, and when MG132 was combined with ACC treatment, the EIL2-GFP signal was at a similar intensity compared to the ACC treatment alone. This result suggest that EIL2 stability is not regulated by the 26S proteasome, and that the increased signal after an ACC treatment is probably due to the transcriptional up-regulation via ethylene signaling. This may also suggest that EIL2 seems to be controlled by a different regulation mechanism than EIN3, which is regulated by proteasomal degradation.

**Figure 5.**
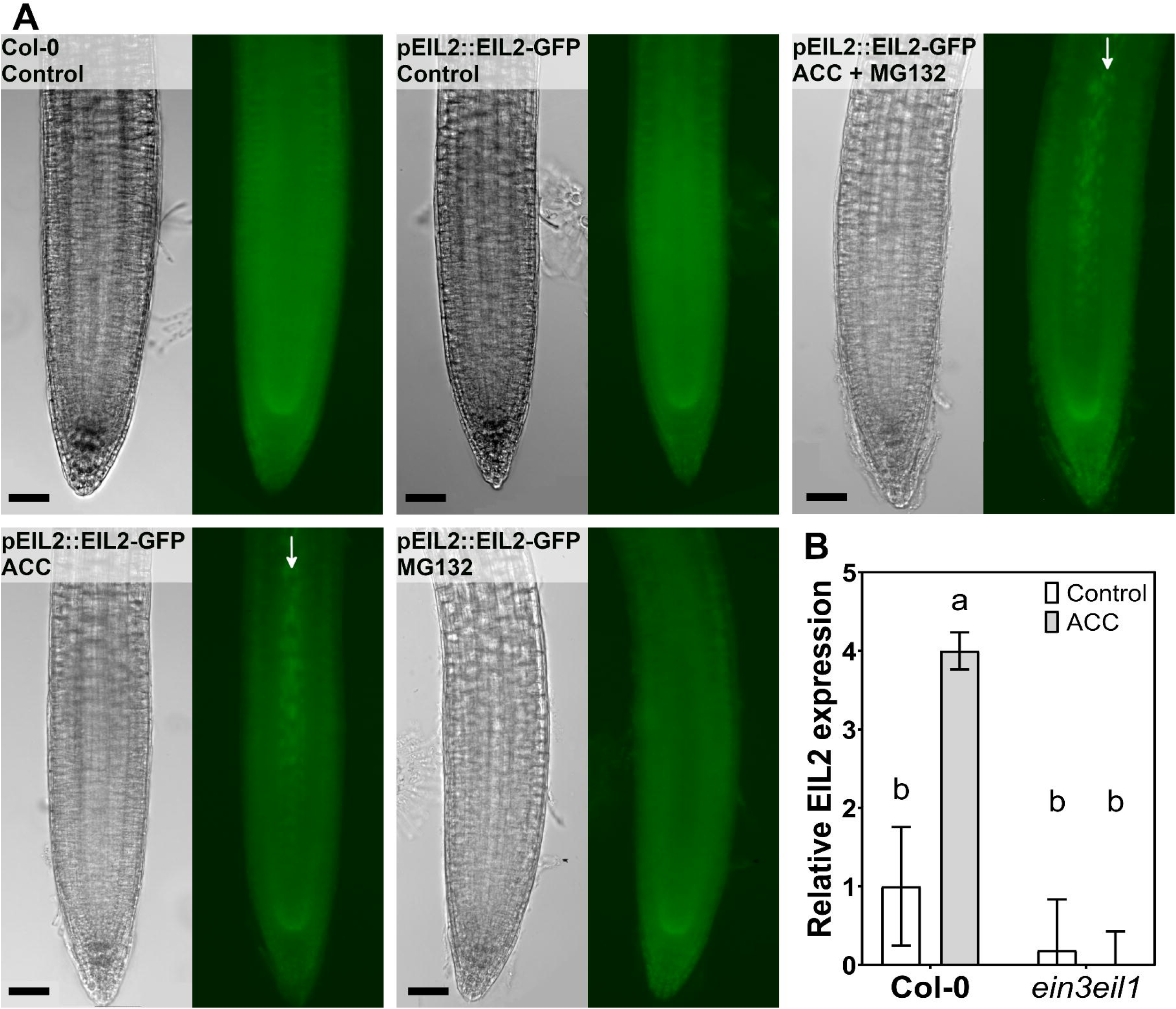
EIL2 signal increases in the stele in response to ACC, but not MG132 treatment. (A) Col-0 and pEIL2::EIL2-GFP translational reporter (C-terminal) were treated for 4 h with 10 μM ACC, 50 μM MG132, ACC + MG132 combined and a control of DMSO only. The EIL2:GFP signal in the stele is indicated by white arrows. Scale bar is 50 μm. (B) Relative expression of EIL2 in 10 day-old light grown seedlings of Col-0 and *the ein3eil1* mutant after 2 μm ACC treatment.

*We* confirmed that EIL2 levels are predominantly transcriptionally regulated by ACC using qPCR (Figure 5B). The native *EIL2* expression was barely detectable, but was significantly enhanced after an ACC treatment in Col-0. We also showed that this ACC-induced expression of *EIL2* is EIN3 and EIN3/EIL1 dependent, because ACC could not stimulate *EIL2* expression in the *ein3* and *ein3eil1* mutants. An analysis of the *EIL2* promotor region also revealed several EIN3 Binding Sequences (EBS) (Supplementary Figure S7) but only 3-4 Kb upstream of the start codon. Although we have showed that *EIL2* expression is ethylene and EIN3/EIL1 dependent, it is not yet clear if EIN3/EIL1 directly controls *EIL2* expression or if other downstream ERF’s are involved in this. Several ERF-specific DNA-binding sites were also retrieved in the promotor of *EIL2* (Supplementary Figure S7).

## Discussion

EIL2, a close homolog of the master ethylene transcription factor EIN3, is considered to be a minor regulator of ethylene signaling, based on the observation that *ein3-1;eil1-3;ebf1-3;ebf2-3* mutants remain ethylene insensitive, despite having a functional EIL2. We now showed that *EIL2* gets expressed upon ethylene signaling in an EIN3/EIL1-dependent way, suggesting EIL2 acts downstream of the first transcriptional wave evoked by EIN3/EIL1^40^. Interestingly, EIL2 seems to be controlled differently compared to EIN3^19^, as EIL2 protein stability is not regulated by proteasomal degradation.

Interestingly, our phylogenetic analysis revealed that EIL2 is a *Brassicaceae*-specific homolog of EIN3, not present in other plant family. This observation suggests EIL2 must have acquired a dedicated function in the *Brassicaceae* family, probably to fine-tune certain ethylene responses. We have now shown that EIL2 plays an important role in several developmental processes, albeit more subtle compared to EIN3/EIL1. We observed that EIL2 controls hypocotyl elongation in the light and is involved in lateral root formation and the regulation of flowering time in *Arabidopsis thaliana*.

### EIL2 promotes hypocotyl elongation and lateral root formation

Ethylene plays an important role in many vegetative developmental processes^2^, yet the role of EIL2 in directing ethylene signaling remained largely unexplored. Our results showed that EIL2 seems to play a role in the ACC-induced hypocotyl elongation response in the light. Previously, this response was attributed to ethylene signaling^41^, and ethylene insensitive mutants (e.g. *etr1-3, ein2-1* and *ein3-1*)displayed a reduced elongation in the light^41^, similar as the *eil2* mutant. Ethylene overproducers (e.g. *eto2*)^42^ and an *EIN3* overexpression^43^ line displayed similar phenotypes in the absence of ethylene. It was shown that hypocotyl elongation is established through the crosstalk action between ethylene (via EIN3 and ERF1) and the Phytochrome Interacting Factors (PIFs), Constitutive Morphogenesis1 (COP1), Long Hypocotyl5 (HY5) complex during seedling photomorphogenesis^42,43^. Perhaps EIN2 is involved in directing hypocotyl elongation via this complex, downstream of EIN3/EIL1.

*Eil2* plants also displayed fewer lateral roots when grown in the light. It was shown before that lateral root development is influenced by ethylene signaling, along with auxin crosstalk^44^. In general, an enhanced ethylene production (ACC feeding or *eto1* mutants)^45,46^ or constitutive ethylene signaling (*ctr1-1* mutants)^45^ lead to fewer lateral roots, while ethylene insensitive mutants (*etr1-3, etr1-1, ein2-5*)^46,47^ display more lateral roots, suggesting ethylene negatively regulates lateral root development. However, other studies have reported conflicting results, namely that ethylene insensitive mutants (*etr1-3*, *ein2-5* and ein3-1eil1-1)^48,49^ do not have more lateral roots. Our results now show that *eil2*, *ein3*and *ein3/eil1* all have fewer lateral roots compared to wild-type plants, grown under control conditions in the absence of ACC, suggesting that EIN3 and its homologs are essential to drive lateral root development. Perhaps EIL2 and/or EIN3/EIL1 are needed to steer the auxin flows that controls lateral root initiation and outgrowth^50^. A possible explanation for the contrasting roles of EIN3/EIL1 and EIL2 in directing lateral root formation might be differences in growth and culture conditions, because it was shown that light^51,52^, nutrients (N, P)^45,49^ and sugars^53^ can all influence lateral root development in crosstalk with ethylene and auxins.

### EIL2 directs flowering time distinctively from EIN3

It was previously established that ethylene is involved in the regulation of flowering time. For example, externally fed ACC leads to delayed flowering *in vitro*^54,55^. Furthermore, ethylene overproducing lines (eřo) show early flowering (1 day)^56^, while some single *acs* (*acs1*, *acs6*, *acs7* and *acs9*) as well as several higher order *acs* mutants, that produce less ethylene, display an early flowering phenotype (up to 6 days)^57^. These findings indicate that ethylene (or ACC) is a negative regulator of flowering time. On the other hand, ethylene insensitive mutants, including *etr1-1*, *ein2-1* and *ein3-1*, display delayed flowering, probably by slowing down the vegetative to reproductive transition^56^. Contradictory, the constitutive ethylene responsive mutant *ctr1-1* also shows a dramatic delay in flowering time (1-2 weeks)^12,54^. Collectively, literature shows that ethylene treatments, ethylene overproduction or constitutive ethylene signaling delay flowering, and surprisingly, a reduced ethylene production or ethylene insensitivity also delay flowering, suggesting there is a dual role for ethylene production and/or signaling to control bolting in Arabidopsis. It was shown that ethylene cross talks with gibberellins (GA) to direct flowering, as the constitutive ethylene signaling mutant *ctr1-1* has lower GA levels, which stabilize DELLA proteins, which are suppressors of the expression of meristem identity genes (*LFY* and SOC1)^54^. The downstream ethylene responsive transcription factor ERF1^54^ and ERF110^55^ are also involved in controlling flowering time. Overexpressing *ERF1* (*35S:ERF1*) drastically delays flowering^54^, and silencing *ERF110* also delays flowering (6 days)^55^. ERF110 gets differentially phosphorylated at serine 62 and is controlled via a EIN2-dependent and EIN2-independent signaling pathways to steer flowering. Our results now show that *eil2* plants flowered faster (only 1 day) suggesting that EIL2 slightly represses flowering. This is in opposition to EIN3, which is a positive regulator of flowering. Perhaps EIL2 regulates flowering time through an interaction with GA signaling, as previously it was shown that EIL2 can interact with the DELLA protein RGA^58^. Our EIL2 findings corroborate that regulation of flowering time by ethylene production and/or signaling is dual and complex, perhaps specific for *Brassicaceae*, and likely influenced through environmental interactions. Ethylene can exert both positive and negative effects on flowering time, and the mechanism how EIL2 takes part in this response, requires further investigation.

## Conclusions

The ethylene transcription factor EIL2 is a close homolog of EIN3 and EIL1, and only retrieved in plants of the *Brassicaceae* family. EIL2 is ethylene responsive and operates downstream of EIN3/EIL1, despite not being controlled by proteasomal degradation. The *EIL2* gene is only weakly expressed in *Arabidopsis thaliana*. EIL2 has some discrete but mild functions in development, as it is involved in ethylene-induced hypocotyl elongation in the light, and it takes part in lateral root development and flowering time control. The transcriptional network and the downstream target genes of EIL2 remain to be revealed.

## Acknowledgements

This work is financially supported by the Research Foundation Flanders (FWO) by a PhD fellowship to MH (SB/1S18717N), a postdoctoral fellowship to WM (1207022N) and research grants to BVDP ( FWO Project G092419N and G0G0219N) and the KU Leuven start-up grant (STGBF/16/005) and C1 research grant (C14/18/056).

## Author contribution

MH performed the GWAS screen. MH, JVH, WM performed the experimental work. BVDP wrote the manuscript and MH, JVH and WM reviewed the manuscript. BVDP supervised the work and provided resources.

## References

1. Abeles F, Morgan P, Saltveit M. Ethylene in Plants. San Diego: Academic Press Inc; 1992.

2. Van de Poel B, Smet D, Van Der Straeten D. Ethylene and Hormonal Cross Talk in Vegetative. Plant Physiol. 2015;169(September):61–72. doi:10.1104/pp.15.00724

3. Ju C, Van De Poel B, Cooper ED, et al. Conservation of ethylene as a plant hormone over 450 million years of evolution. Nat Plants. 2015;1(January):1–7. doi:10.1038/nplants.2014.4

4. Li D, Flores-Sandoval E, Ahtesham U, et al. Ethylene-independent functions of the ethylene precursor ACC in Marchantia polymorpha 2. Nat Plants. 2020;6(11):1335–1344. doi:10.1038/s41477-020-00784-y

5. Li FW, Brouwer P, Carretero-Paulet L, et al. Fern genomes elucidate land plant evolution and cyanobacterial symbioses. Nat Plants. 2018;4(7):460–472. doi:10.1038/s41477-018-0188-8

6. Chen Y, Randlett MD, Findell L, Schaller GE. Localization of the Ethylene Receptor ETR1 to the Endoplasmic Reticulum of Arabidopsis. J Biol Chem. 2002;277(22):19861–19866. doi:10.1074/jbc.M201286200

7. Grefen C, Städele K, Růžička K, Obrdlik P, Harter K, Horák J. Subcellular localization and in vivo interactions of the Arabidopsis thaliana ethylene receptor family members. Mol Plant. 2008;1(2):308–320. doi:10.1093/mp/ssm015

8. Hua J, Meyerowitz EM. Ethylene responses are negatively regulated by a receptor gene family in Arabidopsis thaliana. Cell, 1998;94(2):261–271. doi:10.1016/S0092-8674(00)81425-7

9. Chang C, Kwok SF, Bleecker AB, et al. Arabidopsis Ethylene-Response Gene ETR1: Similarity of Product to Two-Component. Science (80-). 1993;262(5133):539–544.

10. Binder BM. The ethylene receptors⍰: Complex perception for a simple gas. 2008;175:8–17. doi:10.1016/j.plantsci.2007.12.001

11. Clark KL, Larsen PB, Wang X, Chang C. Association of the Arabidopsis CTR1 Raf-like kinase with the ETR1 and ERS ethylene receptors. Proc Natl Acad Sci U S A. 1998;95(9):5401–5406. doi:10.1073/pnas.95.9.5401

12. Kieber JJ, Rothenberg M, Roman G, Feldmann KA, Ecker JR. CTR1, a Negative Regulator of the Ethylene Pathway in Arabidopsis, Encodes a Member of the Raf Family of Protein Kinases. Cell, 1993;72(3):427–441. doi:10.1016/0092-8674(93)90119-B

13. Ju C, Mee G, Marie J, et al. CTR1 phosphorylates the central regulator EIN2 to control ethylene hormone signaling from the ER membrane to the nucleus in Arabidopsis. 2012. doi:10.1073/pnas.1214848109/-/DCSupplemental. www.pnas.org/cgi/doi/10.1073/pnas.1214848109

14. Alonso JM, Hirayama T, Roman G, Nourizadeh S, Ecker JR. EIN2, a Bifunctional Transducer of Ethylene and Stress Responses in *Arabidopsis*. Science (80-). 1999;284(5423):2148–2152. doi:10.1126/science.284.5423.2148

15. Roman G, Lubarsky B, Kieber JJ, Rothenberg M, Ecker R. Genetic Analysis of Ethylene Signal Transduction in Arabidopsis thaliana: Five Novel Mutant Loci Integrated into a Stress Response Pathway. Genetics. 1995;139:1393–1409.

16. Qiao H, Chang KN, Yazaki J, Ecker JR. Interplay between ethylene, ETP1/ETP2 F-box proteins, and degradation of EIN2 triggers ethylene responses in Arabidopsis. Genes Dev. 2009;23(4):512–521. doi:10.1101/gad.1765709

17. Qiao H, Shen Z, Huang SSC, et al. Processing and subcellular trafficking of ER-tethered EIN2 control response to ethylene gas. Science (80-). 2012;338(6105):390–393. doi:10.H26/science.1225974

18. Wen X, Zhang C, Ji Y, et al. Activation of ethylene signaling is mediated by nuclear translocation of the cleaved EIN2 carboxyl terminus. Cell Res. 2012;22:1613–1616. doi:10.1038/cr.2012.145

19. Guo H, Ecker JR. Plant responses to ethylene gas are mediated by SCFEBF1/EBF2-dependent proteolysis of EIN3 transcription factor. Cell. 2003;115(6):667–677. doi:10.1016/S0092-8674(03)00969-3

20. Potuschak T, Lechner E, Parmentier Y, et al. EIN3-dependent regulation of plant ethylene hormone signaling by two Arabidopsis F box proteins: EBF1 and EBF2. Cell. 2003;115(6):679–689. doi:10.1016/S0092-8674(03)00968-1

21. Gagne JM, Smalle J, Gingerich DJ, et al. Arabidopsis EIN3-binding F-box 1 and 2 form ubiquitin-protein ligases that repress ethylene action and promote growth by directing EIN3 degradation. Proc Natl Acad Sci. 2004;101(17):6803–6808.

22. Merchante C, Brumos J, Yun J, et al. Gene-Specific Translation Regulation Mediated by the Hormone-Signaling Molecule EIN2. Cell. 2015;163(3):684–697. doi:10.1016/j.cell.2015.09.036

23. Li W, Ma M, Feng Y, et al. EIN2-Directed Translational Regulation of Ethylene Signaling in Arabidopsis Article EIN2-Directed Translational Regulation of Ethylene Signaling in Arabidopsis. Cell. 163(3):670–683. doi:10.1016/j.cell.2015.09.037

24. Solano R, Stepanova A, Chao Q, Ecker JR. Nuclear events in ethylene signaling: A transcriptional cascade mediated by ETHYLENE-INSENSITIVE3 and ETHYLENE-RESPONSE-FACTOR1. Genes Dev. 1998;12(23):3703–3714. doi:10.1101/gad.12.23.3703

25. Song J, Zhu C, Zhang X, et al. Biochemical and structural insights into the mechanism of DNA recognition by arabidopsis ETHYLENE INSENSITIVE3. PLoS One. 2015;10(9). doi:10.1371/journal.pone.0137439

26. Zhang F, Qi B, Wang L, et al. EIN2-dependent regulation of acetylation of histone H3K14 and non-canonical histone H3K23 in ethylene signalling. Nat Commun. 2016;7:1–14. doi:10.1038/ncomms13018

27. Wang L, Zhang Z, Zhang F, et al. EIN2-directed histone acetylation requires EIN3-mediated positive feedback regulation in response to ethylene. Plant Cell. 2021;33(2):322–337. doi:10.1093/plcell/koaa029

28. Chao Q, Rothenberg M, Solano R, Roman G, Terzaghi W, Ecker JR. Activation of the ethylene gas response pathway in Arabidopsis by the nuclear protein ETHYLENE-INSENSITIVE3 and related proteins. Cell. 1997;89(7):H33–1144. doi:10.1016/S0092-8674(00)80300-1

29. Binder BM, Walker JM, Gagne JM, et al. The Arabidopsis EIN3 binding F-box proteins EBF1 and EBF2 have distinct but overlapping roles in ethylene signaling. Plant Cell. 2007;19(2):509–523. doi:10.1105/tpc.106.048140

30. Dolgikh VA, Pukhovaya EM, Zemlyanskaya E V. Shaping Ethylene Response: The Role of EIN3/EI11 Transcription Factors. Front Plant Sci. 2019;10(August):1–9. doi:10.3389/fpls.2019.01030

31. Maruyama-nakashita A, Nakamura Y, Tohge T, Saito K, Takahashi H. Arabidopsis SLIM1 Is a Central Transcriptional Regulator of Plant Sulfur Response and Metabolism. 2006;18(November):3235–3251. doi:10.1105/tpc.106.046458

32. Wawrzynska A, Sirko A. To control and to be controlled: understanding the Arabidopsis SLIM1 function in sulfur deficiency through comprehensive investigation of the EIL protein family. Front Plant Sci. 2014;5(October):1–7. doi:10.3389/fpls.2014.00575

33. Smyth DR, Bowman JL, Meyerowitz EM. Early Flower Development in Arabidopsis. Plant Cell. 1990;2(August):755–767. papers3://publication/uuid/0BFEAF50-2E5A-458F-B90B-DFF364DFB25D.

34. Seren Ü, Vilhjálmsson BJ, Horton MW, et al. GWAPP: A web application for genome-Wide association mapping in arabidopsis. Plant Cell. 2013;24(12):4793–4805. doi:10.1105/tpc.112.108068

35. Letunic I, Bork P. Interactive tree of life (iTOL) v3: an online tool for the display and annotation of phylogenetic and other trees. Nucleic Acids Res. 2016;44(W1):W242–W245. doi:10.1093/nar/gkw290

36. Boavida LC, McCormick S. Temperature as a determinant factor for increased and reproducible in vitro pollen germination in Arabidopsis thaliana. Plant J. 2007;52(3):570–582. doi:10.1111/j.1365-313X.2007.03248.x

37. Mori T, Kuroiwa H, Higashiyama T, Kuroiwa T. GENERATIVE CELL SPECIFIC 1 is essential for angiosperm fertilization. Nat Cell Biol. 2006;8(1):64–71. doi:10.1038/ncb1345

38. Waese J, Fan J, Pasha A, et al. ePlantØ: Visualizing and Exploring Multiple Levels of Data for Hypothesis Generation in Plant Biology. Plant Cell. 2017;29(August):1806–1821. doi:10.1105/tpc.17.00073

39. Feng Y, Xu P, Li B, et al. Ethylene promotes root hair growth through coordinated EIN3/EI11 and RHD6/RS11 activity in Arabidopsis. Proc Natl Acad Sci U S A. 2017;114(52):13834–13839. doi:10.1073/pnas.1711723115

40. Chang KN, Zhong S, Weirauch MT, et al. Temporal transcriptional response to ethylene gas drives growth hormone cross-regulation in Arabidopsis. Elife. 2013;2013(2):1–20. doi:10.7554/eLife.00675

41. Smalle J, Haegman M, Kurepa J, Van Montagu M M Van, Straeten D V. Ethylene can stimulate Arabidopsis hypocotyl elongation in the light. Proc Natl Acad Sci U S A. 1997;94(6):2756–2761. doi:10.1073/pnas.94.6.2756

42. Yu Y, Wang J, Zhang Z, et al. Ethylene Promotes Hypocotyl Growth and HY5 Degradation by Enhancing the Movement of COP1 to the Nucleus in the Light. PLoS Genet. 2013;9(12). doi:10.1371/journal.pgen.1004025

43. Zhong S, Shi H, Xue C, et al. A molecular framework of light-controlled phytohormone action in arabidopsis. Curr Biol. 2012;22(16):1530–1535. doi:10.1016/j.cub.2012.06.039

44. Lewis DR, Negi S, Sukumar P, Muday GK. Ethylene inhibits lateral root development, increases IAA transport and expression of PIN3 and PIN7 auxin efflux carriers. Development. 2011;138(16):3485–3495. doi:10.1242/dev.065102

45. Lopez-Bucio J, Hernandez-Abreu E, Sanchez-Calderon L, Nieto-Jacobo M, Simpson J, Herrera-Estrella L. Phosphate availability alters architecture and causes changes in hormone sensitivity in the arabidopsis root system. Plant Physiol. 2002;129:244–256.

46. Negi S, Ivanchenko MG, Muday GK. Ethylene regulates lateral root formation and auxin transport in Arabidopsis thaliana. Plant J. 2008;55(2):175–187. doi:10.llll/j.1365-313X.2008.03495.x

47. Feigl G, Horváth E, Molnár Á, Oláh D, Poór P, Kolbert Z. Ethylene-Nitric Oxide Interplay During Selenium-induced Lateral Root Emergence in Arabidopsis. J Plant Growth Regul. 2019;38(4):1481–1488. doi:10.1007/s00344-019-09950-9

48. Abozeid A, Ying Z, Lin Y, Liu J, Zhang Z, Zhonghua T. Ethylene improves root system development under cadmium stress by modulating superoxide anion concnetration in Arabidopsis thaliana. Front Plant Sci. 2017;8(253).

49. Li G, Li B, Dong G, Feng X, Kronzucker HJ, Shi W. Ammonium-induced shoot ethylene production is associated with the inhibition of lateral root formation in Arabidopsis. J Exp Bot. 2013;64(5):1413–1425. doi:10.1093/jxb/ert019

50. Muday GK, Rahman A, Binder BM. Auxin and ethylene: Collaborators or competitors? Trends Plant Sci. 2012;17(4):181–195. doi:10.1016/j.tplants.2012.02.001

51. Zeng J, Wang Q, Lin J, Deng K, Zhao X. Arabidopsis cryptochrome-1 restrains lateral roots growth by inhibiting auxin transport. J Plant Physiol. 2010;167:670–673. doi:10.1016/j.jplph.2009.12.003

52. Gelderen K Van, Kang C, Li P, Pierik R. Regulation of Lateral Root Development by Shoot-Sensed Far-Red Light via HY5 Is Nitrate-Dependent and Involves the NRT2.1 Nitrate Transporter. Front Plant Sci. 2021;12(March):1–12. doi:10.3389/fpls.2021.660870

53. Singh M, Gupta A, Laxmi A. Ethylene acts as a negative regulator of glucose induced lateral root emergence in Arabidopsis. Plant Signal Behav. 2015;10:9. doi:10.1080/15592324.2015.1058460

54. Achard P, Baghour M, Chapple A, et al. The plant stress hormone ethylene controls floral transition via DELLA-dependent regulation of floral meristem-identity genes. Proc Natl Acad Sci USA. 2007;104(15):6484–6489. doi:10.1073/pnas.0610717104

55. Zhu L, Liu D, Li Y, Li N. Functional phosphoproteomic analysis reveals that serine-62-phosphorylated isoform of ethylene response factorllO is involved in Arabidopsis bolting. Plant Physiol. 2013;161:904–917.

56. Ogawara T, Higashi K, Kamada H, Ezura H. Ethylene advances the transition from vegetative growth to flowering in Arabidopsis thaliana. J Plant Physiol. 2003;160(H):1335–1340. doi:10.1078/0176-1617-01129

57. Tsuchisaka A, Yu G, Jin H, et al. A combinatorial interplay among the 1-aminocyclopropane-1-carboxylate isoforms regulates ethylene biosynthesis in Arabidopsis thaliana. Genetics. 2009;183(3):979–1003. doi:10.1534/genetics.109.107102

58. An F, Zhang X, Zhu Z, et al. Coordinated regulation of apical hook development by gibberellins and ethylene in etiolated Arabidopsis seedlings. Cell Res. 2012;22(5):915–927. doi:10.1038/cr.2012.29

